# Alternate response of *Bacillus anthracis atxA* and *acpA* to serum, HCO_3_^-^ and CO_2_

**DOI:** 10.1101/2022.02.21.481328

**Authors:** Itai Glinert, Elad Bar-David, Amir Ben-Shmuel, Assa Sittner, Reut Puni, Shira Laredo, David Kobiler, Shay Weiss, Haim Levy

## Abstract

*Bacillus anthracis* overcomes host immune responses by producing capsule and secreting toxins. Production of these virulence factors in response to entering the host environment was shown to be regulated by *atxA,* the major virulence regulator, known to be activated by HCO_3_^-^ and CO_2_. While toxin production is regulated directly by *atxA,* capsule production is independently mediated by two regulators; *acpA* and *acpB.* In addition, it was demonstrated that *acpA* has at least two promotors, one of them shared with *atxA.* We used a genetic approach to study capsule and toxin production under different conditions. Unlike previous works utilizing NBY-HCO_3_^-^ medium under CO_2_ enriched conditions, we used a sDMEM-based medium. Thus, toxin and capsule production can be induced in ambient or CO_2_ enriched atmosphere. Using this system, we could differentiate between induction by 10% NRS, 10% CO_2_ or 0.75% HCO_3_^-^. In response to high CO_2_, capsule production is induced by *acpA* in an *atxA*-independent manner, with little to no toxin (protective antigen PA) production. *atxA* is activated in response to serum independently of CO_2_, inducing toxin and capsule production in an *acpA* or *acpB* dependent manner. HCO_3_^-^ was also found to activate *atxA*, but in non-physiological concentrations. Our findings may help explain the first stages of inhalational infection, in which spores germinating in dendritic cells require protection (by encapsulation) without affecting cell migration to the draining lymph-node by toxin secretion.

## Introduction

*Bacillus anthracis,* the causative agent of anthrax, is a gram positive, spore forming bacterium that infects humans via three major routes; inhalational (lung), cutaneous (skin) and gastrointestinal (gut) [1, 2]. The infectious form of *B. anthracis* is the spore, durable and environmentally stable for decades. In response to entering the host, spores germinate and overcome the immune system utilizing two major virulence factors; the poly-δ-D-glutamic antiphagocytic capsule and the tripartite toxin system [3]. Capsule producing genes are encoded as a polycistronic gene cluster on the pXO2 virulence plasmid, comprised of the *capB,C,A,D,E* genes. CapB and CapC are the D-glutamic acid polymerization enzymes, while CapA and CapE form the transport channel that exports the polymer from the cytoplasm through the cell wall and membrane to the cell surface [4–6]. Once there, CapD hydrolyzes the polymer to shorter chains and covalently links them to the cell wall [7]. CapD also regulates the length of the bound polymer chains through its hydrolysis activity [4, 8, 9]. The tripartite toxins are encoded by the *pag* (protective antigen-PA), *lef* (lethal factor-LF) and *cya* (edema factor-EF) genes, located on the pXO1 virulence plasmid [1]. LF is a metalloprotease that specifically cleaves members of the MAP kinase regulatory pathway of mammalian cells [10]. EF is a potent calmudolin dependent adenylate cyclase that interferes with cell regulation by elevating the internal cAMP levels [11]. Both LF and EF are driven into mammalian cells by PA, which binds specific receptors ANTXR1 (TEM8) and ANTXR2 (CMG2) [12]. Following binding to the receptor, PA is processed by cell-associated furin, activating oligomerization, forming a heptamer that binds LF and EF [12, 13]. The complex is then phagocyted and upon lysosomal fusion, the pH drop causes a conformational shift resulting in LF and EF injection into the cytoplasm. These toxin moieties then disrupt cell activity by disrupting intra-cellular signaling. Intoxication of immune cells results in their inactivation, disrupting host immune responses [12].

The AtxA was shown to be the major virulence regular of *B. anthracis* [2, 14]. This pXO1-located gene (encoding for the AtxA protein) activates a cascade of regulatory processes, resulting in up and down regulation of both chromosomal and plasmid encoded genes [15]. Capsule production is regulated by two pXO2 encoded genes – *acpA* [15] and *acpB* [16, 17]. Though it was demonstrated that AtxA regulates these two genes [18], it is not essential for capsule production since ΔpXO1 strains remain capable of capsule production [19, 20]. Since *B. anthracis* does not produce toxins or capsule under normal laboratory growth conditions, specific host-simulating growth conditions were developed. It was reported that growth of *B. anthracis* in NBY broth supplemented with glucose and bicarbonate in 5-15% CO_2_ atmosphere induces toxin secretion and capsule production [16, 20–22]. These growth conditions were used to study different aspects of virulence regulation in *B. anthracis.*

Resulting findings indicated that AtxA in induced in response to HCO_3_^-^ and CO_2_, activating both toxin and capsule production. This concept implies that toxins and capsule are produced simultaneously in the host.

Previously we reported that growth of *B. anthracis* in supplemented DMEM (a high glucose cell culture medium supplemented with pyruvate, glutamine, nonessential amino acids, (henceforth sDMEM) with the addition of 10% serum, and incubated in a 10% CO_2_ atmosphere, induces virulence factor production [17, 19]. These conditions were used to examine the effects of serum (here - normal rabbit serum, NRS), HCO_3_^-^ and/or CO_2_ atmosphere, on the regulation of toxin and capsule production. This was coupled with a systematic genetic approach, to dissect the regulation of the bacterial virulence response. Unlike previous reports, we show that *AtxA* is induced by NRS or HCO_3_^-^, but not CO_2_. The capsule regulator AcpA can be induced by CO_2_ in an *atxA*-independent manner, or by NRS in an *atxA-dependent* manner. AcpB was activated only in an *atxA*-dependent manner, with *atxA* deletion resulting in complete abrogation of *acpB* capsule regulation. Our results indicate that bacteria is capable of independent capsule production, uncoupled from toxin production, but that toxins are always be co-induced with the capsule.

## Materials and Methods

### Bacterial strains, media and growth conditions

Bacterial strains used in this study are listed in **Table 1**. For the induction of toxins and capsule production, we employed a modified DMEM (supplemented with 4mM L-glutamine, 1 mM Sodium pyruvate, 1% non-essential amino acid) that was supplemented with 10% normal rabbit serum or 0.75% NaHCO3 (Biological Industries – Israel) – hence sDMEM. Spores of the different mutants were seeded at a concentration of 5×10^5^ CFU/ml and grown in 96 well tissue culture plates (100μl per well) for 5 or 24h at 37oC in ambient or 10% CO2 atmosphere.

**Table 1:** Bacterial strains used in this study

| Strain | Description/characteristics | Source |
| --- | --- | --- |
| **<u>B. anthracis</u>** |  |  |
| Vollum | ATCC 14578 | IIBR collection |
| VollumΔpXO1 | Complete deletion of the plasmid pXO1 | IIBR collection |
| VollumΔatxA | Complete deletion of the atxA gene | This study |
| VollumΔacpA | Complete deletion of the acpA gene | [17] |
| VollumΔacpB | Complete deletion of the acpB gene | [17] |
| VollumΔacpAΔacpB | Complete deletion of the acpA and acpB genes | [17] |
| VollumΔpagΔcyaΔlef (ΔTox) | Complete deletion of the pag, lef and cya genes | [23] |
| VollumΔpagΔcyaΔlefΔacpA | Complete deletion of the acpA gene in the VollumΔpagΔcyaΔlef mutant | [17] |
| VollumΔpagΔcyaΔlefΔacpB | Complete deletion of the acpB gene in the VollumΔpagΔcyaΔlef mutant | [17] |
| VollumΔpagΔcyaΔlefΔacpAΔacpB | Complete deletion of the acpA and acpB genes in the VollumΔpagΔcyaΔlef mutant | [17] |
| VollumΔpagΔcyaΔlefΔacpAΔatxA | Complete deletion of the atxA gene in the VollumΔpagΔcyaΔlefΔacpA mutant | This study |
| VollumΔpagΔcyaΔlefΔacpBΔatxA | Complete deletion of the atxA gene in the VollumΔpagΔcyaΔlefΔacpB mutant | This study |
| VollumΔpXO2 ΔbclA::<br>pag<sub>prom</sub>::capA-E | Genomic insertion of the pagA promoter in front of the CAP operon replacing the bclA gene in the VollumΔpXO2 | [17] |
| VollumΔpXO2 ΔbclA::<br>pag<sub>prom</sub>::capA-EΔatxA | Deletion of the atxA gene from the strain having Genomic insertion of the pagA promoter in front of the CAP operon replacing the bclA gene in the VollumΔpXO2 | [17] |

### Mutant strain construction

Oligonucleotide primers used in this study were previously described [17, 19, 23]. The oligonucleotide primers were designed according to the genomic sequence of *B. anthracis* Ames strain. Genomic DNA (containing the chromosomal DNA and the native plasmids, pX01 and pX02) for cloning the target gene fragments was extracted from the Vollum strain as previously described [24]. Target genes were disrupted by homologous recombination, using a previously described method [25]. In general, gene deletion or insertion was accomplished by a marker-less allelic exchange technique that replaced the complete coding region with the SpeI restriction site or the desired sequence. At the end of the procedure the resulting mutants did not code for any foreign sequences and the only modification is the desired gene insertion or deletion. Deletion of the atxA gene was performed as previously described [25]. All the mutants were tested for their ability to produce capsule by incubation in sDMEM. The capsule was visualized by negative staining with India ink.

### Toxin quantification

Protective antigen (PA) concentration was determine by capture ELISA using the combination of a polyclonal and monoclonal αPA antibodies previously described [23]

## Results

### The effect of supplements and growth condition on capsule production in sDMEM medium

The basic medium used to test capsule and toxin induction was high glucose DMEM supplanted with glutamine, pyruvate and non-essential amino acids, hence sDMEM. We first tested the effect of growth media volume on capsule production in a 96-well tissue culture microplate format. This was done to allow higher-throughput testing of growth conditions and genetic manipulations. We inoculated *B. anthracis* Vollum spores (5×10^5^ CFU/ml) into 100μl, 200μl and 300μl sDMEM at 37°C in ambient atmosphere for 24h. No capsule production could be detected in 100μl sDMEM (**Figure 1**). However, increasing the culture volume resulted in partial (200μl) or full (300μl) capsule production (**Figure 1**). These findings could be possibly explained by higher CO_2_ concentrations reached in the well when bacteria are grown in larger volumes, due to a lower surface area to volume ratio, limiting gas exchange. This is coupled with a larger volume of multiplying and respiring bacteria, further increasing CO_2_ concentrations. This hypothesis is supported by the finding that growth in 10% CO_2_ atmosphere does induce capsule production in 100μl sDMEM (**Figure 1**). Having established non-capsule inducing growth conditions for our experiments, we next tested the effect of adding normal rabbit serum (NRS) to growth media. NRS was shown to induce capsule production, regardless of CO_2_ concentrations in the incubator’s atmosphere, with all the bacteria grown with NRS encapsulated in ambient or 10% CO_2_ atmosphere (**Figure 1**).

**Figure 1.**
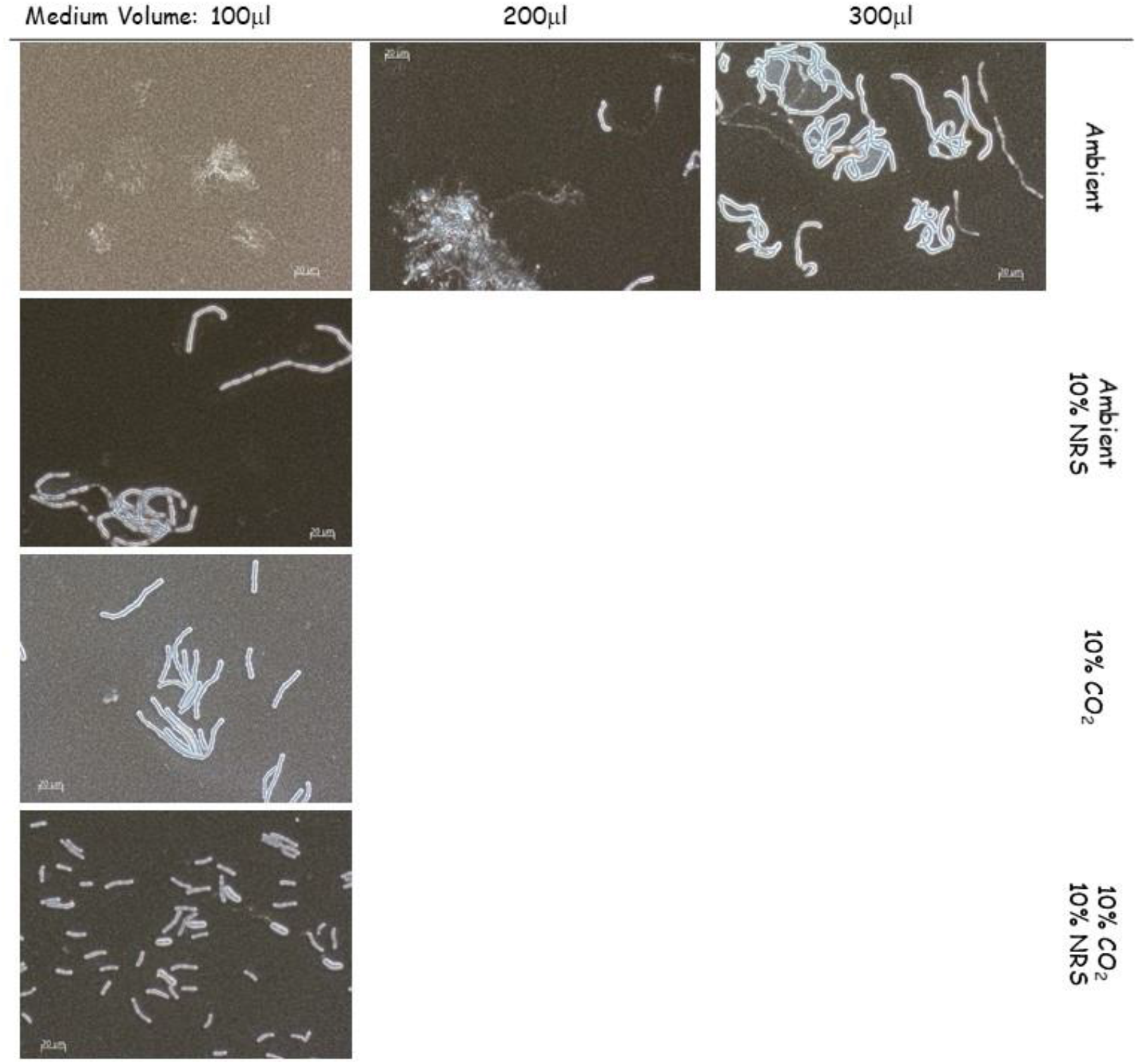
The effect of growth conditions on capsule induction by *B. anthracis* Vollum. Spores were seeded into 100μl, 200μl or 300μl of sDMEM and incubated at 37°C in an ambient atmosphere (upper panel) for 24h. To examine the effect of serum or CO_2_ on capsule production, spores were seeded into 100μl sDMEM or sDMED-NRS (supplemented with 10% NRS). Samples were incubated at 37°C in under ambient or 10% CO_2_ atmosphere (see right hand side legend) for 24h. Capsule was imaged by India ink negative staining (capsule presence forms a typical bright outer layer).

### The role of *atxA* on capsule production and toxin secretion under different growth conditions

To determine the role of *atxA* on capsule and toxin production, we compared capsule production of the wild type Vollum strain with the ΔpXO1 or *ΔatxA* mutants (**Table 1**) following growth in sDMEM under the different growth conditions (NRS/CO_2_ presence). None of the strains produced capsule under ambient atmosphere in un-supplemented sDMEM (**Figure 2**). Addition of 10% NRS, under ambient atmosphere, induced capsule production by the wild type Vollum strain but not by the *atxA* null mutants (both ΔpXO1 or *Δat×A,* **Figure 2**). Under 10% CO_2_, all three strains produced capsule. We therefore conclude that the response to NRS is *atxA* dependent, as lacking either the atxA gene or the entire pXO1 plasmid abrogates this response (**Figure 2**). These results also demonstrate the presence of another, atxA independent mechanism, responding to CO_2_. However, the effect of NRS in the presence of CO_2_ on capsule production under these conditions could not be resolved.

**Figure 2.**
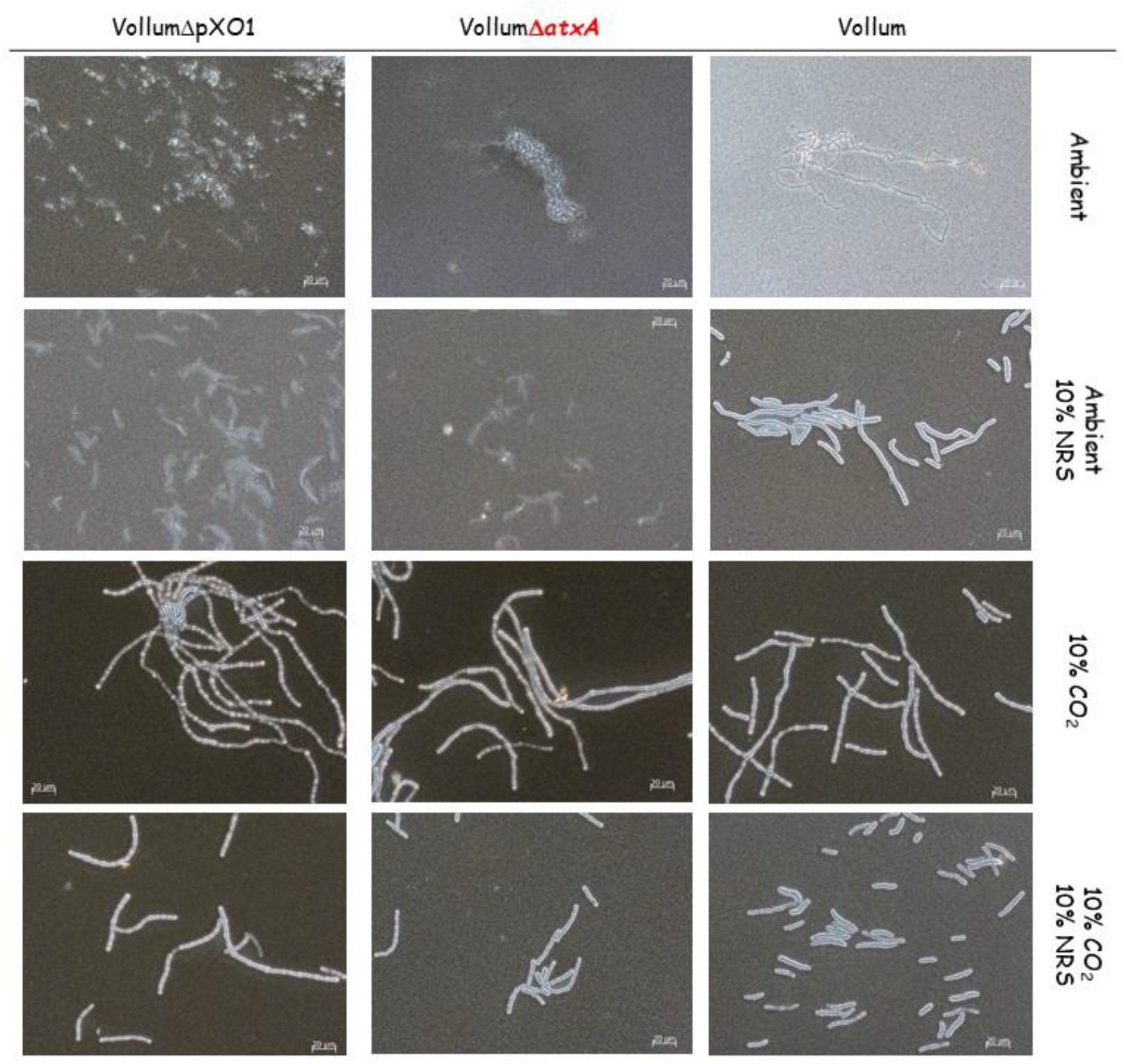
Capsule production of the Vollum wild type, ΔpXO1 or *ΔatXA* mutants under different growth conditions. Spores of the wild type and different mutants (top panel) were seeded into 100μl of sDMEM. The same strains were grown in sDMEM supplemented with 10% NRS (second row). The same conditions were applied in the presence of 10% CO_2_ (lower two rows) for 24h. Capsule was imaged by India ink negative staining (capsule presence forms a typical bright outer layer).

As *atxA,* the major virulence regulator, regulates toxin expression (LF, EF and PA), we tested their secretion into growth medium after growth, both for the Vollum and the VollumΔ*atxA* mutant (the ΔpXO1 mutant lacks the entire set of toxin genes and is therefore irrelevant for such a test). Toxins secretion was determined by ELISA for the most abundant component, PA. For the VollumΔ*atxA* mutant, PA was undetectable, both under 10% CO_2_, with 10% NRS and in the presence of both. These results confirm the role of atxA in toxin expression regulation (**Table 2**). For the wild type Vollum, no toxins were induced in sDMEM alone coupled with under ambient atmosphere. The addition of 10% NRS alone induced high PA expression. sDMEM without NRS under 10% CO_2_ again did not induce PA expression (**Table 2**). However, the addition of both NRS and CO_2_ resulted in reduced PA expression (about a 70% reduction) (**Table 2**). This finding was maintained throughout different, independent experiments. However, we cannot exclude the possibility that this difference resulted from growth rates variation under the tested growth conditions. No PA secretion could be detected in the *atxA* null mutant under any of the tested growth conditions. Our finding that 10% CO_2_ atmosphere in itself does not activate *atxA*-dependent toxin expression is surprising, since it was repeatedly demonstrated such activation is achievable by HCO_3_^-^ addition. We therefore tested both capsule production and toxin expression in sDMEM supplemented with 0.75% HCO_3_^-^ under 10% CO_2_ atmosphere. These atmospheric conditions were chosen since HCO_3_^-^ in aqueous media is in equilibrium with CO_2_ and H_2_O, dramatically increasing the levels of the soluble CO_2_ in the medium even under ambient atmosphere, and we sought to expose our experimental controls to conditions as similar as possible.

**Table 2.**
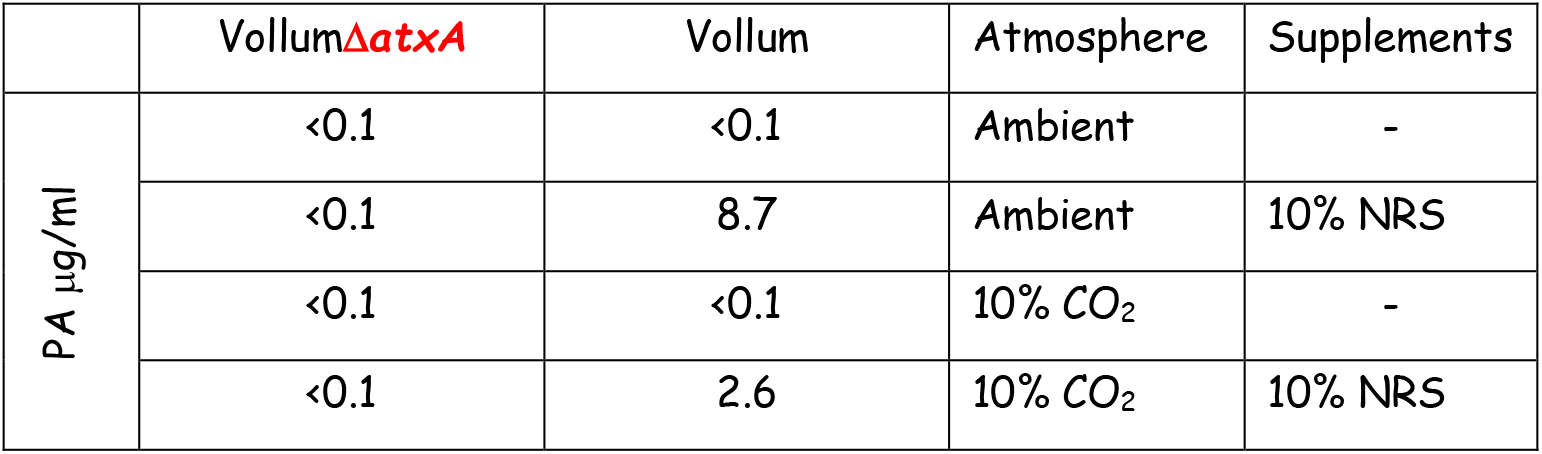
Protective Antigen secretion of Vollum or VollumΔ*atxA* under different growth conditions.

According to our previous findings, capsule production is CO_2_ dependent. Therefore, we found that the above tested strains were encapsulated under the conditions applied (**Figure 3**). PA secretion was tested only for Vollum and VollumΔ*atxA*. Supplementing the sDMEM with 0.75% HCO_3_^-^ induced PA secretion by the Vollum strain but not by the VollumΔ*atxA* mutant. This indicates that HCO_3_^-^ induces toxin secretion in an *atxA* dependent manner (**Table 3**). To validate these findings, we used a previously reported VollumΔpXO2 chimera in which we substituted the genomic *bclA* gene (which is a redundant exosporium glycoprotein) with a CAP operon altered to be regulated by the PA_prom_ (**Table 1**). Thus, this mutant serves here as an indicator for AtxA dependent activation of the PA promotor, which results in capsule production (assayable readout). This mutant was then used as the basis for another mutant strain, in which the *atxA* gene was deleted, on top of the existing genetic alterations (**Table 1**). Thus, the exact role *atxA* plays in the regulation on the PA-promotor under the tested conditions could be resolved.

**Figure 3.**
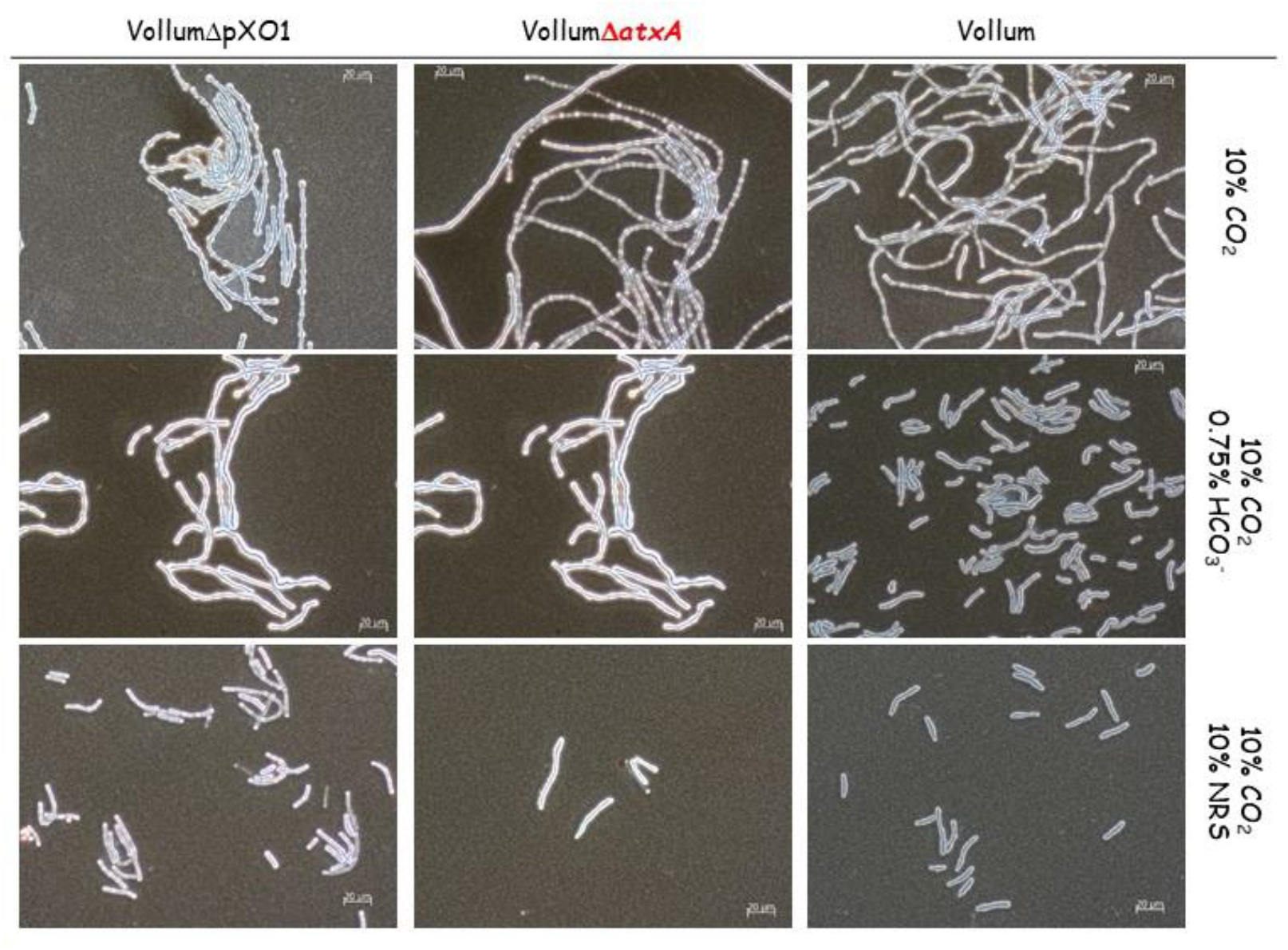
Effect of 0.75% HCO_3_^-^ on capsule production of Vollum, VollumΔ*atxA* or VollumΔpXO1. Spores were seeded into 100μl of sDMEM as is or supplemented with 0.75% HCO_3_^-^ or 10%NRS, and incubated at 37°C under 10% CO_2_ for 24h. Capsule was imaged by India ink negative staining (capsule presence forms a typical bright outer layer).The different mutations are indicated in the top panel.

**Table 3.** PA secretion by Vollum and VollumΔ*atxA* in sDMEM supplemented with HCO_3_^-^.

| | Vollum$\Delta\text{atxA}$ | Vollum | Atmosphere | Supplements |
| --- | --- | --- | --- | --- |
| PA $\mu\text{g/ml}$ | <0.1 | <0.1 | 10% $\text{CO}_2$ | - |
| | <0.1 | 3.12 | 10% $\text{CO}_2$ | 0.75% $\text{HCO}_3^-$ |
| | <0.1 | 5.3 | 10% $\text{CO}_2$ | 10% NRS |

While 10% CO_2_ atmosphere induced capsule production in the Vollum wild type strain, little or no capsule production could be detected in the VollumΔpXO2Δ*bclA*::*pagA_Prom_*-capA-E chimeric strain (**Figure 4**). Supplementing sDMEM with 0.75% HCO_3_^-^ resulted in capsule production in the Vollum and VollumΔpXO2Δ*bclA*::*pagA_prom_*-capA-E chimera. However, deletion of the *atxA* gene, in the background of the chimera mutant described, resulted in loss of capsule production (VollumΔpXO2Δ*bclA*::*pagA_prom_*-capA-*EΔatxA,* **Figure 4**). Similar results were obtained by supplementing sDMEM with 10% NRS. These results complement the PA secretion experiment, indicating that HCO_3_^-^ and NRS induce toxin secretion in an *atxA* dependent manner.

**Figure 4.**
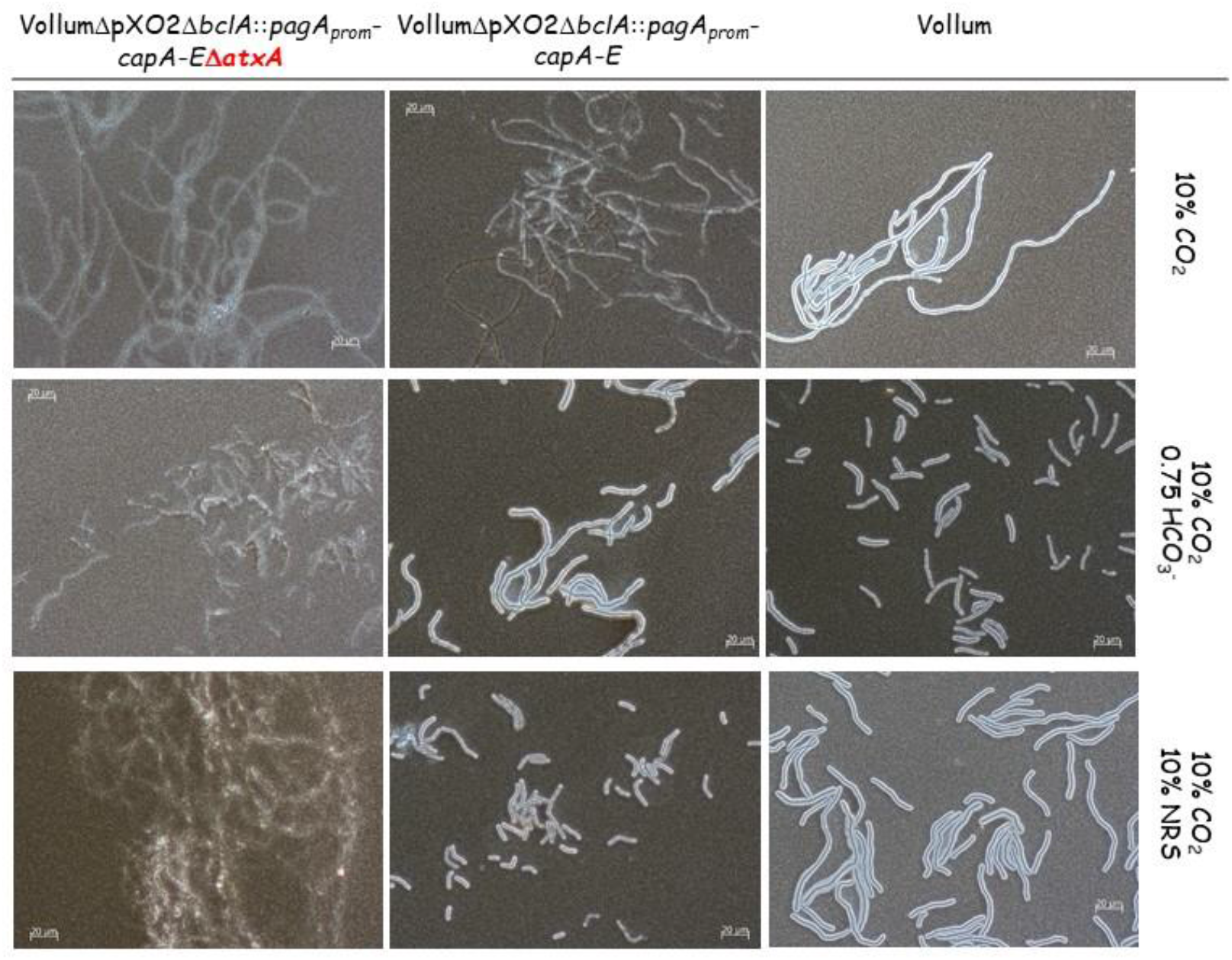
Capsule production in response to HCO_3_^-^ in a PAprom-regulated genomic CAP operon. Spores were seeded into 100μl of sDMEM as is or supplemented with 0.75% HCO_3_^-^ or 10% NRS and incubated at 37°C under an atmosphere of 10% CO_2_ for 24h. Capsule was imaged by India ink negative staining (capsule presence forms a typical bright outer layer).The different mutations are indicated (top panel).

### The effect of short incubations on capsule and toxins production in response the different growth conditions

Aerobic growth affects different parameters of the liquid medium such as pH and O_2_/CO_2_ concentrations, especially when the bacteria reaches high concentration (CFU/ml). As was shown previously [26], capsule production and toxin secretion can be detected as early as 2-5h of growth in sDMEM under 10% CO_2_ atmosphere. To minimize changes in media conditions resulting from bacterial growth, we examined capsule production and PA secretion after 5h growth in different growth conditions. Vollum growth under ambient atmosphere did not result in any capsule accumulation or toxin secretion following 24h incubation (**Figure 2**, **Table 2**). This was true also for a short (5h) incubation (**Figure 5**, **Table 4**). Supplementing the media with 10% NRS induced capsule production and PA secretion following 24h incubation under ambient atmosphere (**Figure 2**, **Table 2**). A shorter (5h) incubation induced measurable PA secretion (**Table 4**) but no capsule production (**Figure 5**). This PA secretion is AtxA dependent, as deletion of the *atxA* gene resulted in no PA accumulation following 5h (**Table 4**) or 24h incubation (**Table 2**). Incubating the bacteria in 10% CO_2_ atmosphere for 5h resulted in capsule production (**Figure 5**), similarly to that observed following a 24h incubation (**Figures 1-3**). This capsule accumulation was AtxA independent and was not significantly affected by the addition of 10% NRS or HCO_3_^-^ (**Figure 5**). PA secretion was not induced by 10% CO_2_ in itself, but required addition of NRS or HCO_3_^-^ (**Table 4**), however unlike the 24h incubation (**Table 3**), the amount of PA secreted in response to HCO_3_^-^ was significantly less (~10%) than of that induced by NRS (**Table 4**). Under all conditions the PA secretion was AtxA dependent.

**Figure 5.**
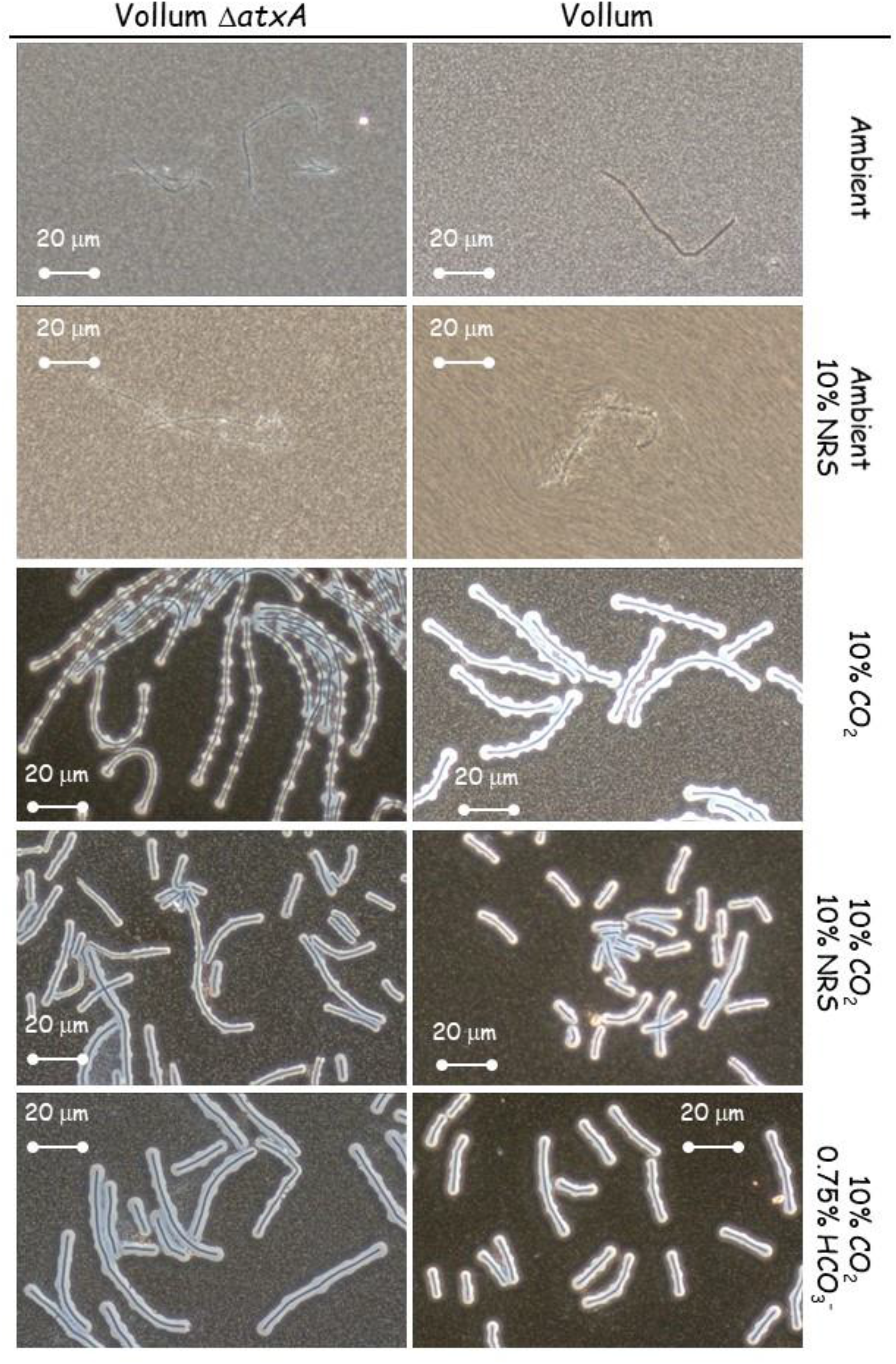
Effect of deleting *atxA* on capsule production after a short (5h) incubation under different growth conditions. Spores of the wild type and Δ*atxA* mutant (top panel) were seeded into 100μl of sDMEM as is or supplemented with 10% NRS and incubated at 37°C in an ambient or 10% CO_2_ atmosphere (as indicated on the right) for 24h. Capsule was imaged by India ink negative staining (capsule presence forms a typical bright outer layer).

**Table 4.** PA secretion by Vollum and VollumΔ*atxA* following 5h growth in different conditions

| | VollumΔatxA | Vollum | Atmosphere | Supplements |
| --- | --- | --- | --- | --- |
| PA μg/ml | <0.1 | <0.1 | Ambient | - |
|  | <0.1 | 1.89 | Ambient | 10% NRS |
|  | <0.1 | <0.1 | 10% CO<sub>2</sub> | - |
|  | - | 4.25 | 10% CO<sub>2</sub> | 10% NRS |
|  | - | 0.29 | 10% CO<sub>2</sub> | 0.75% HCO<sub>3</sub><sup>-</sup> |

### Regulation of *acpA* and *acpB* in response to different growth conditions

Toxin production is induced in an *atxA* dependent manner in response to HCO_3_^-^ or NRS, while capsule production is also induced by a CO_2_ enriched (10%) atmosphere. Capsule production is known to be regulated by two regulatory proteins, AcpA and AcpB. We therefore tested the effect of deleting each of these genes on capsule production in response to different growth conditions. The complete coding region of *acpA* or *acpB* was deleted independently in the background of wild type Vollum or the toxin deficient mutant - VollumΔTox (VollumΔ*pag*Δ*cya*Δ*lef* **Table 1**). As was previously shown for the wild type Vollum strain (**Figure 2**), none of these mutants produced capsule following growth in sDMEM under ambient atmosphere (**Figure 6**). The presence of either *acpA* or *acpB* is sufficient for capsule production in sDMEM supplemented with 10% NRS, regardless to the presence or absence of 10% CO_2_ atmosphere (**Figure 6**). However, in the absence of NRS, only AcpA expressing mutants (lacking *acpB)* produce significant capsule when grown in 10% CO_2_ atmosphere. Adding 0.75% HCO_3_^-^ to sDMEM induced capsule production in the presence of either *acpA* or *acpB.* Mutants lacking *acpA* (expressing only *acpB*), did not produce significant capsule in 10% CO_2_ atmosphere. To examine the role of *atxA* in these processes, we deleted the *atxA* gene in the background of our VollumΔToxΔ*acpA* or *ΔacpB* mutants. Deleting *atxA* in the VollumΔToxΔ*acpA*, expressing *acpB,* abolished capsule production under all tested conditions (**Figure 7**). However, deleting *atxA* in the VollumΔToxΔ*acpB*, expressing *acpA,* did not affect capsule accumulation, compared to the AtxA expressing mutant. This finding indicates that *acpA* operates in an AtxA independent manner (**Figure 7**). As we demonstrated (**Figure 2**), capsule production could be induced in ambient atmosphere by adding 10% NRS to sDMEM. This induction is AtxA dependent, since no capsule production was detected in the VollumΔ*atxA* mutant under these conditions (**Figure 2**). Since AcpA dependent capsule production in 10% CO_2_ atmosphere was AtxA independent, we tested the role of AtxA on AcpA dependent capsule production in response to 10% NRS in ambient atmosphere. As 10% NRS induced capsule production of VollumΔ*acpB* under ambient atmosphere (**Figure 6, Figure 8**) we tested the effect of *atxA* deletion on capsule production under these conditions. Unlike the CO_2_ induction, under ambient atmosphere, AcpA dependent capsule production in response to NRS is AtxA dependent (**Figure 8**).

**Figure 6.**
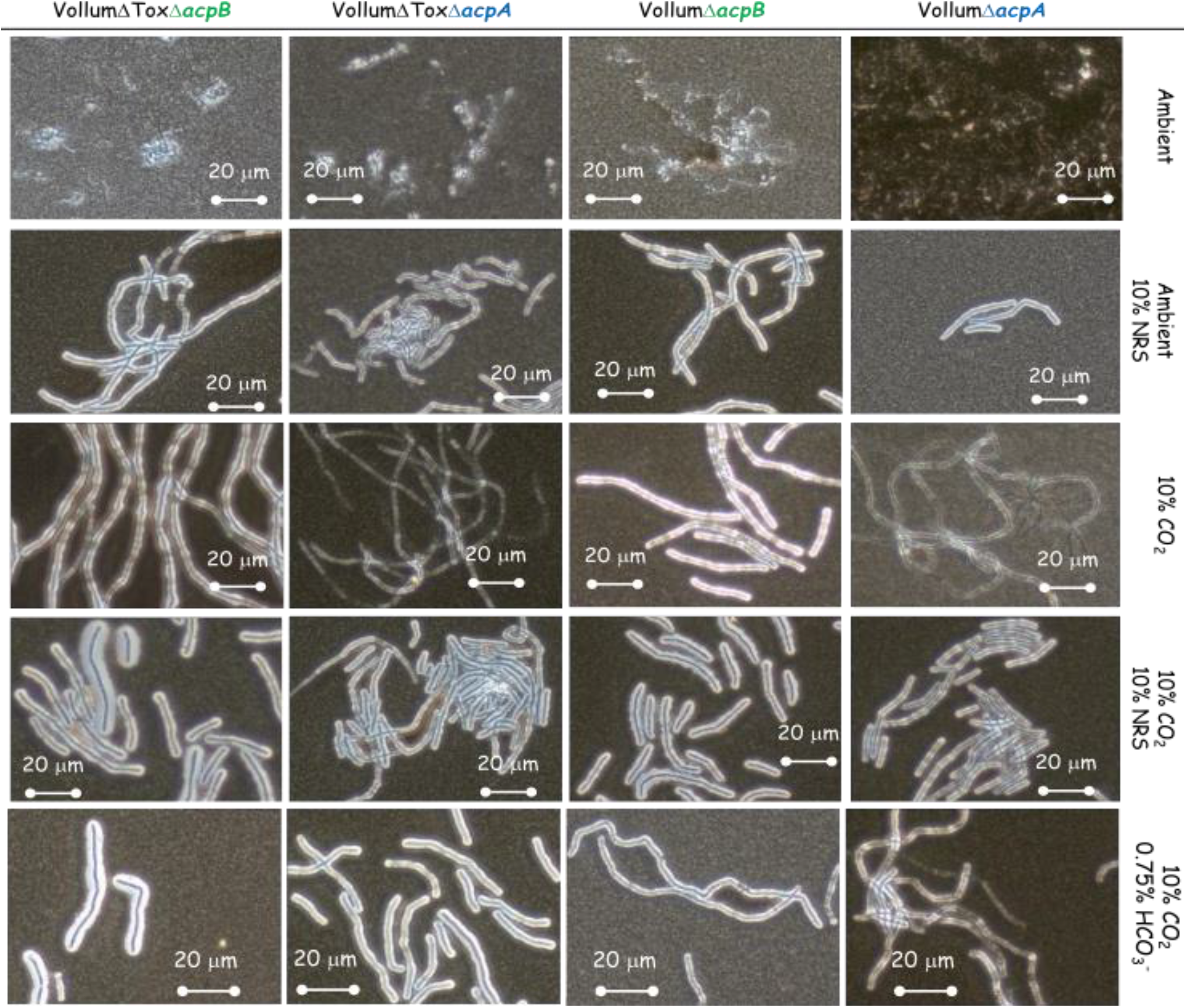
The effect of absence of *acpA* or *acpB* on capsule production in response to 10% NRS under ambient or 10% CO_2_ atmosphere. Spores of the *ΔacpA* or *acpB* mutants (top panel) were seeded into 100μl of sDMEM as is or supplemented with0.75% HCO_3_^-^ or 10% NRS and incubated at 37°C in an ambient or 10% CO_2_ atmosphere (as indicated on the right) for 24h. Capsule was imaged by India ink negative staining (capsule presence forms a typical bright outer layer).

**Figure 7.**
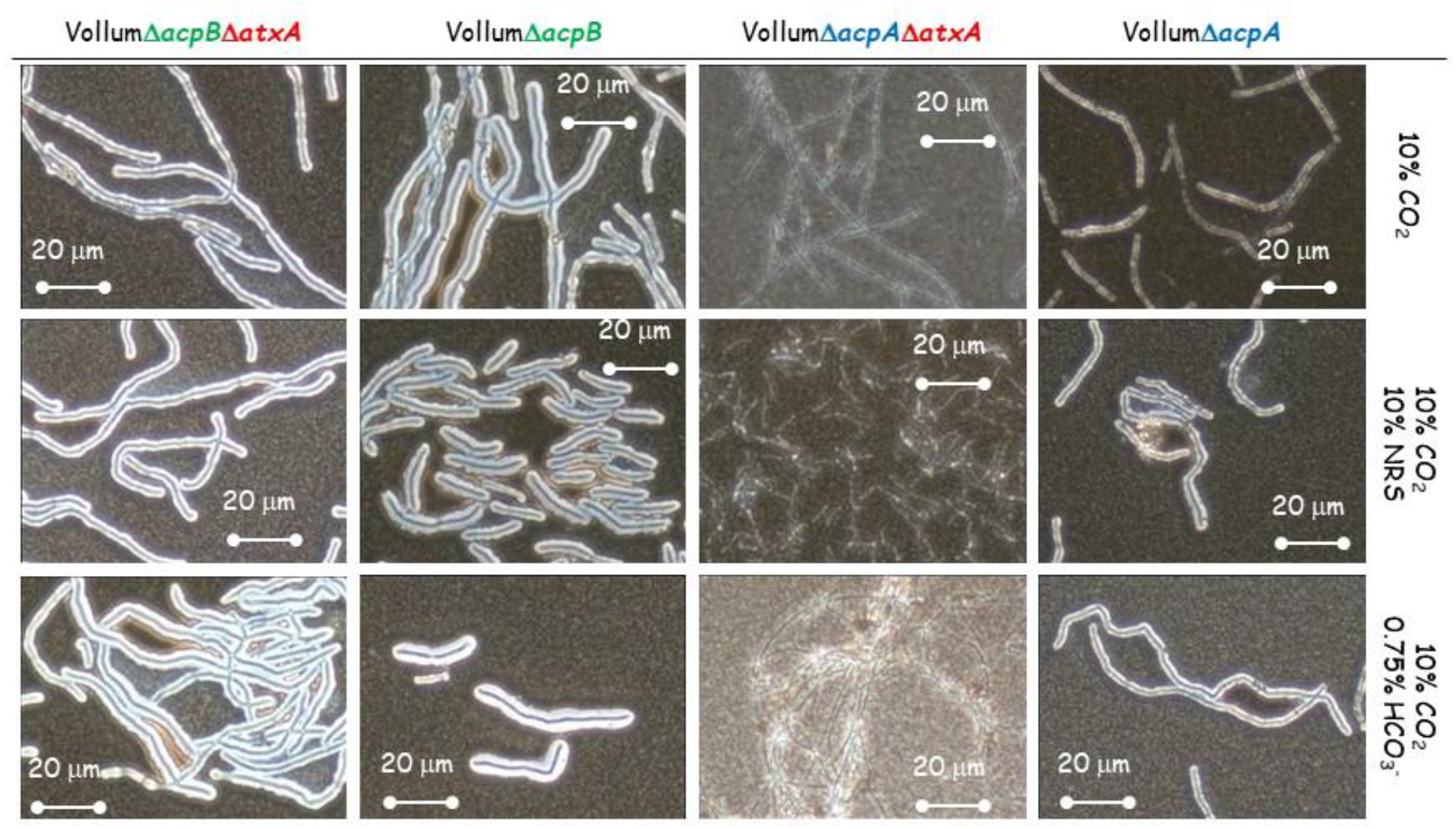
Effect of AtxA on capsule production in the presence of either AcpA or AcpB. Spores of the different mutants (top panel) were seeded into 100μl of sDMEM as is or supplemented with 0.75% HCO_3_^-^ or 10% NRS and incubated at 37°C in 10% CO_2_ atmosphere (as indicated on the right) for 24h. Capsule was imaged by India ink negative staining (capsule presence forms a typical bright outer layer).

**Figure 8.**
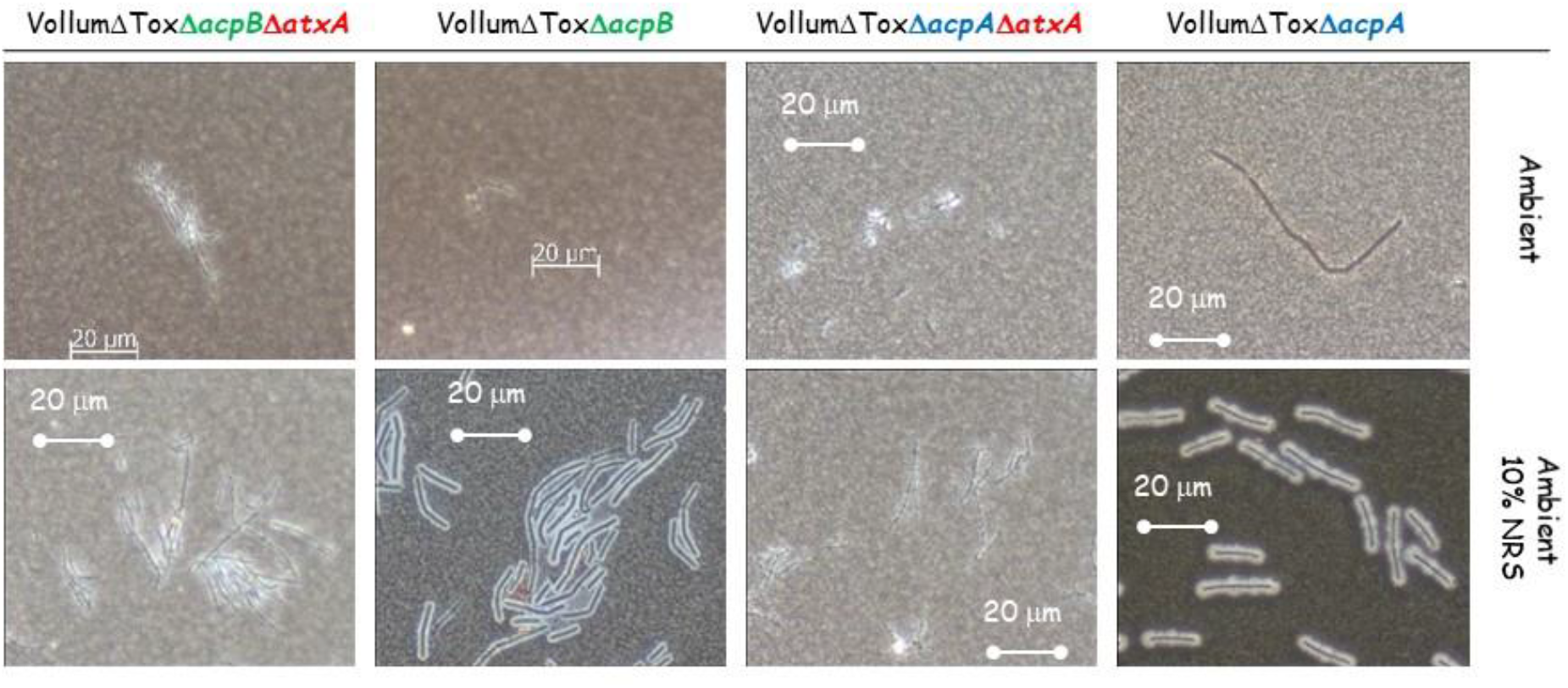
Effect of AtxA on capsule production in response to 10% NRS in ambient atmosphere. Spores of the different mutants (top panel) were seeded into 100μl of sDMEM as is or supplemented with 10% NRS and incubated at 37°C in ambient atmosphere (as indicated on the right) for 24h. Capsule was imaged by India ink negative staining (capsule presence forms a typical bright outer layer).

### Gas content of the supplemented and basic DMEM media under the different growth conditions

Gas content was determined by Abbott iSTAT blood gas analyzer. The different media were incubated at 37°C in ambient or 10% CO_2_ atmosphere for 4 h and analyzed using the EC8+ cartridge. The results shown in **Table 5** indicate that the main parameter that is affected by the 10% CO_2_ is the soluble CO_2_ (PCO_2_) rather than HCO_3_^-^. Unlike the HCO_3_^-^ levels that are higher under all growth conditions from those considered normal for human blood, the PCO_2_ levels in the presence of 10%CO_2_ are well within or very close to normal values.

**Table 5.** Gas analysis of the sDMEM media under the different growth conditions

| 10% CO<sub>2</sub> | - | + | + | - | + | Normal<br>arterial | Normal<br>venous |
| --- | --- | --- | --- | --- | --- | --- | --- |
| Supplement | - | - | 10% FCS | 10% FCS | 0.75% HCO<sub>3</sub><br>1:1* |  |  |
| pH | 8.0 | 7.691 | 7.447 | 8.149 | 7.972 | 7.35-7.45 | 7.31-7.41 |
| PCO<sub>2</sub> (mmHg) | 18.4 | 34.9 | 58.1 | 14.2 | 34.3 | 35-45 | 41-51 |
| PO<sub>2</sub> (mmHg) | 230 | 148 | 126 | 173 | 183 | 80-100 | 35-40 |
| HCO<sub>3</sub> (mmol/L) | 48.7 | 42.2 | 40.1 | 49.5 | 79.2 | 22-26 | 22-26 |
| sO<sub>2</sub> (%) | 100 | 100 | 99 | 100 | 100 | 95-100 | / |

\*at the end of the 4h incubation the media was diluted 1:1 in sDMEM prior to the analysis since the undiluted media was off scale.

## Discussion

Successful invasion requires the pathogen regulate its virulence factors in a way that will maximize their effect on host defense mechanisms. The trigger for such activation is usually host derived. This can be biological (such as proteins) of physical (pH or temperature for example). *B. anthracis* naturally infects humans following spore inhalation, contact with broken skin or ingestion of undercook contaminated meat. These routes present different environmental conditions [2]. It was previously demonstrated that toxin secretion and capsule production could be induced by growing the bacteria in culture media supplemented with HCO_3_^-^ or serum (10-50%) in a CO_2_ enriched (5-15%) atmosphere [5, 16, 18, 19, 27–29]. HCO_3_^-^ /CO_2_ condition were commonly used to study *atxA, acpA* and *acpB* regulation and their effect on toxin and capsule biosynthetic genes [16, 18, 30]. Since these conditions always included these two components, it was concluded that *atxA* was induced in response to CO_2_, regulating the induction of *acpA* and *acpB.* Although in some reports, capsule production was shown to be AtxA dependent [16, 31], the fact that ΔpXO1 variants are encapsulated contradicts this finding, alluding to additional, AtxA independent regulation of the process [19, 27]. The use of sDMEM as growth media enabled the examination of the effect of CO_2_, HCO_3_^-^ and serum on these processes.

The parameter of soluble CO_2_ is influenced by multiple parameters, such as surface area to volume ratio and aerobic bacterial growth. Therefore, normal growth conditions were set to 100μl media/well (96 well tissue culture plate) for 24h at 37°C under ambient atmosphere (**Figure 1**). This baseline enabled testing the effect of different supplements and/or growth conditions on capsule (**Figure 1**) or toxin (**Figure 2, Table 2**) induction. Capsule production is induced by the addition of 10% NRS or growth under a 10% CO_2_ atmosphere (**Figure 1**). The serum capsule induction (under ambient atmosphere) is *atxA* dependent (**Figure 2**), since there was no significant capsule production in mutants that do not express AtxA (VollumΔ*atxA* and VollumΔpXO1). Alternatively, capsule production in response to CO_2_ enriched atmosphere is AtxA independent, as there is no significant difference in capsule production, under these conditions, between AtxA expressing and *atxA* null mutants (**Figures 2, 3**). Toxin secretion, as determined by measuring PA media concentrations, is serum dependent (**Tables 2, 3**), as PA could be detected only in NRS supplemented sDMEM regardless of CO_2_ enrichment. Adding HCO3^-^ induced toxin secretion in an *atxA* dependent manner, similarly to serum (**Figure 3, Table 3**), with PA expressed by the wild type Vollum and not by the *atxA* null mutant. The same NRS/HCO_3_^-^ dependence and CO_2_ independence of PA induction was demonstrated using a previously described mutant strain, in which pXO2 is missing and a copy of the CAP operon was inserted chromosomally, regulated by a PA promotor [17]. This mutant strain produces capsule when grown in sDMEM supplemented with NRS or HCO_3_^-^ but not in non-supplemented sDMEM under 10% CO_2_ atmosphere.

We also found differences in the speed *B. anthracis* reacts to the different stimuli. This was done by examining toxin secretion and capsule production after a short incubation (5h). HCO_3_^-^ was not as robust as NRS in inducing toxin secretion. Examining PA concertations after 5h incubation in sDMEM supplemented with 0.75% HCO_3_^-^ revealed about 1/10 of the concentration compared to that measured after 24h incubation (**Figure 5, Table 4**). However, NRS induction yielded similar concentrations at these two timepoints. Testing the effect of NRS on capsule production, reveals that 5h incubation under ambient atmosphere, inducing significant PA secretion, does not result in significant capsule production. Growth under a 10% CO_2_ atmosphere induced capsule production at 5h even in the absence of supplemented NRS or HCO_3_^-^ (**Table 4**). Hence, Serum seemed more effective then HCO_3_^-^ in inducing toxin secretion, with two process being AtxA dependent. The AtxA independent induction of capsule production by 10% CO_2_ appeared more effective then AtxA dependent serum induction (**Figure 5**).

Two major regulators; AcpA and AcpB control capsule biosynthesis by promoting transcription of *acpB,C,A,D,E* operon. *acpA* was shown to be regulated by AtxA (activated by NRS), while also depicting an additional *atxA* independent activation capacity by CO_2_ and HCO_3_^-^). This independent activation could be direct or mediated by an as yet unknown, additional promotor, in turn responding to an CO_2_ ingredient. We found that deleting *acpA* causes the bacteria to produce significantly less capsule in response to CO_2_, while maintaining its ability to respond to NRS or HCO_3_^-^ (**Figure 6**). Deletion of *acpB* did not have any effect on capsule production under all tested conditions (**Figure 6**), supporting our previous *in vivo* data, which showed no effect on virulence [17]. A double deletion of *atxA* and *acpA* or *acpB* revealed that AcpB activity is strictly AtxA dependent under all the conditions tested (**Figure7**). AcpA activity is not affected by the absence of *atxA* in the presence of CO_2_ (**Figure7**) but is nulled in response to NRS under ambient atmosphere (**Figure 8**).

Our findings delineate the following regulation cascade; CO_2_ induces capsule production by activating of *acpA* in an AtxA independent manner. Serum activates the AtxA dependent cascade, inducing toxin secretion and eventually capsule production, by activating *acpA* and *acpB* (**Figure 9**). The order of the processes can be deduced from the lack of capsule production following 5h growth in NRS supplemented sDMEM under ambient atmosphere (**Figure 5**). HCO_3_^-^ induces toxin secretion through the AtxA cascade, but in a less efficient manner (compared to NRS, **Table 4**). Direct activation of capsule production by HCO_3_^-^ in an AtxA independent manner could not be eliminated, since even under ambient atmosphere, it modifies the levels of soluble CO_2_ (PCO_2_) and possibly induces capsule production via AcpA (**Table 5**). In terms of the initial infection stages of pathology, this differential regulation of toxins and capsule has a logical role. Inhalational and cutaneous infections involve spore phagocytosis by local innate immune cells and their migration to a draining lymph node. While en route (and within the phagocytic cell), the spore needs to germinate and produce the protective capsule. During this stage, toxin production could prove counterproductive, as it disrupts normal cellular function, possibly arresting the cell and preventing it from reaching the lymph node. Once there, toxin production is desirable, possibly enhancing bacterial release from the cells in the lymph node, to the blood stream, promoting pathogenesis. The pathway sensing serum and CO_2_ remains to be elucidated and requires more research. Such a pathway may prove common to other pathogens as well as possibly providing additional therapeutic targets for intervention.

**Figure 9.**
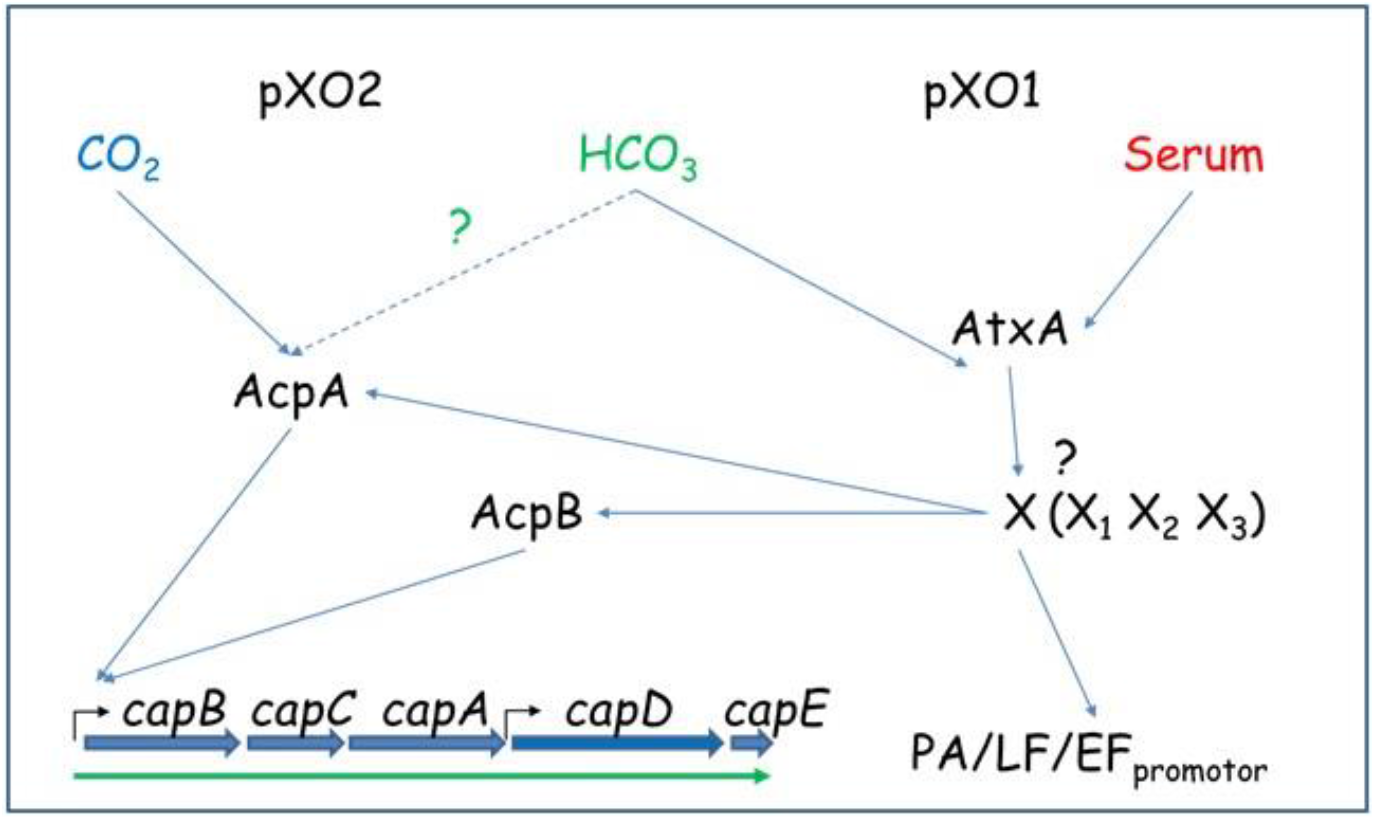
Proposed regulatory scheme for CO_2_, Serum and HCO_3_^-^ regulation of capsule production and toxin secretion.

